# Electron microscopic and crystallographic studies of bacteriophage Sf6 procapsid-like particles assembled from heterologously expressed capsid protein gp5

**DOI:** 10.1101/2023.06.28.546888

**Authors:** Haiyan Zhao, Liang Tang

**Affiliations:** Molecular Biosciences, University of Kansas, 1200 Sunnyside Ave., Lawrence, KS, 66045, USA; Department of Biochemistry and Molecular Genetics, University of Colorado Anschutz Medical School, Aurora, CO, 80045, USA

**Author notes:** **Synopsis** The bacteriophage Sf6 procapsid-like particles were assembled from heterologously expressed capsid protein gp5 and crystallized. Self-rotation function showed the icosahedral symmetry and located the particle in the crystal unit cell.

## Abstract

Many double-stranded DNA (dsDNA) viruses undergo a capsid maturation process during assembly of infectious virus particles, which involves transformation of a metastable capsid precursor called procapsid into a stable, DNA-filled capsid usually with a larger size and a more angular shape. Sf6 is a tailed dsDNA bacteriophage that infects *Shigella flexneri*. The phage Sf6 capsid protein gp5 was heterologously expressed and purified. Electron microscopy showed that the gp5 spontaneously assembled into spherical, procapsid-like particles. We also observed tube-like and cone-shaped particles reminiscent of human immunodeficiency virus. The gp5 procapsid-like particles were crystallized and crystals diffracted beyond 4.3 Å resolution. X-ray data at 5.9 Å resolution were collected with a completeness of 31.1% and an overall *R*merge of 15.0%. The crystals belong to the space group *C*2 with unit cell dimensions of a=973.326 Å, b=568.234 Å, c=565.567 Å, and β=120.540°. Self-rotation function showed the 532 symmetry, confirming formation of icosahedral particles. The particle was situated at the origin of the crystal unit cell with the icosahedral 2-fold axis coinciding with the crystallographic *b* axis, and there is a half of the icosahedral particle in the crystallographic asymmetric unit.

## 1. Introduction

A hallmark in assembly of tailed double-stranded DNA (dsDNA) bacteriophages and herpesviruses is maturation of a metastable capsid precursor called procapsid into the stable, DNA-filled capsid (Veesler & Johnson, 2012), driven by the viral DNA-packaging machinery (Sun *et al*., 2008, Zhao *et al*., 2013, Dai *et al*., 2013). The procapsid is usually smaller and spherical, while the mature capsid exhibits a more angular shape and a larger size. The procapsid-to-capsid maturation process is accompanied with massive structural rearrangement as shown in phages and herpesviruses in the context of isolated particles (Veesler & Johnson, 2012) and in infected cells (Dai *et al*., 2013). The capsid proteins that undergo such massive structural transitions in those viruses share a common “HK97” fold (Suhanovsky & Teschke, 2015, Zhou *et al*., 2014). Structural basis of the procapsid-to-capsid maturation at the atomic detail has been available for phage HK97 (Wikoff *et al*., 2000, Gertsman *et al*., 2009). Recent electron cryo-microscopic structures of phage T7 DNA-free procapsid, mature capsid and a DNA-packaging intermediate shed light on the maturation process at the near atomic level (Guo *et al*., 2014).

Bacteriophage Sf6 belongs to the *Podoviridae* family of tailed dsDNA bacteriophages (Casjens *et al*., 2004, Gemski *et al*., 1975). Sf6 infects *Shigella flexneri*, an important human pathogen that causes acute diarrhea and bacillary dysentery. Sf6 can alter the host’s serotype and virulence by changing the structure of the host lipopolysaccharide through horizontal gene transfer (Lindberg *et al*., 1978). Sf6 is closely related to phages P22, HK620 and ST64T (Casjens *et al*., 2004, Parent *et al*., 2012). The capsid proteins of these phages contain an I-domain in addition to the canonical “HK97” fold (Rizzo *et al*., 2014, Parent *et al*., 2012), and this domain has been shown by mutagenesis studies to be important for folding and stability of capsid proteins and be involved in appropriate assembly of the *T*=7 icosahedral procapsid and capsid (Suhanovsky & Teschke, 2015). Such I-domain is also present in podoviruses CUS-3 (Parent *et al*., 2014) and C1 (Aksyuk *et al*., 2012) as well as in phage T4, a member of the *Myoviridae* (Fokine *et al*., 2005). However, a structure at atomic or near atomic resolution is not available for phage procapsid or capsid with I-domain-containing capsid proteins, hampering understanding of the structural basis for the role of the I-domain in the virus assembly and the procapsid-to-capsid maturation.

We have assembled procapsid-like particles (PLPs) *in vitro* using the heterologously expressed and purified capsid protein gp5 of phage Sf6, and hereby report crystallographic analysis of the gp5 PLPs.

## 2. Materials and methods

### 2.1. Macromolecule production

The coding sequence for the Sf6 capsid protein gp5 was amplified by PCR from the phage Sf6 genomic DNA with the use of the following primers: 5’-CCA GAA TTC ATG CCT AAC AAT CTC GAC -3’, which contains a EcoR1 site followed by the gp5 coding sequence and 5’-GGC CTC GAG CGG ATT ACC GAA GAA CTG-3’, which introduces an Xho1 site followed by a His6 tag after the C-terminus of gp5. The PCR product was digested with the appropriated restriction enzymes, gel purified, and cloned into the same sites of the vector pET28b (Novagen). The plasmid of gp5 was transformed in the *E. coli* strain BL21(DE3) (Novagen). Cells were grown at 30ºC in LB media supplemented with 30 µg/ml Kanamycin until OD600 reached around 0.6, after which IPTG was added to a final concentration of 1mM, and cells were grown for additional 3 hours. Cells were harvested by centrifugation for 10min at 5000 rpm. Cell pellets were resuspended in buffer A (20mM Tris-HCl pH 8.0, 50mM NaCl and 10mM β-mercaptoethanol, 1mg/ml lysozyme and 0.1mg/ml DNaseI) and frozen at -20ºC. Cells were thawed at room temperature, lysed by incubating for 2 hours at the room temperature with gently shaking and centrifuged at 8,000 rpm with a Sorvall SS-34 rotor for 20 minutes at 4°C. The supernatant was transferred to a clean centrifugation tube and centrifuged at 18,000 rpm for 4 hours. The pellets were resuspended in buffer B (20mM Tris-HCl pH 8.0, 50mM NaCl and 1mM EDTA) for 20 hours. The milky gp5 solution was briefly centrifuged at 5000 rpm for 10 minutes. The supernatant was carefully transferred to a clean tube. The final gp5 concentration was 13.4mg/ml in buffer B and was stored at 4 ºC.

The purified gp5 PLPs was diluted by 10-fold immediately prior to adsorption onto an EM grid, followed by negative staining with 1% uranyl acetate using standard procedures as described (van Heel *et al*., 2000). The grid was air dried and examined on an FEI Tecnai F20 transmission electron microscope operating at 200kV.

### 2.2. Crystallization

Crystal screening was originally performed using an in-house set of screening conditions designed for virus crystallization, which contained low concentrations of polyethylene glycol. Conditions that generated crystals were selected for optimization. The final crystals were obtained by hanging drop vapour diffusion, in which 1ul protein was mixed with 1ul well solution containing 5% polyethylene glycol 8000, 100 mM Tris-HCl pH 8.5 and 1M NaCl. Crystals appeared after a few days and grew to 0.4×0.2×0.1 mm3 within two weeks. Crystals were flash-frozen in the well solution containing 30% ethylene glycol prior to X-ray data collection.

### 2.3. Data collection and processing

X-ray data were collected at a cryogenic temperature of 100K on the beamlines 23-ID-B and 23-ID-D at the Advanced Photon Source (APS), using a MAR 300 CCD detector and a Pilatus3 6M detector, respectively. A number of crystals were screened with the exposure time in the range of 2-8 seconds per frame and crystal-to-detector distances of 850-1000 mm. The data used for calculation of self-rotation function were collected at the APS beamline 23-ID-B. A total of 131 frames were collected from a single crystal with an oscillation range of 1°, an exposure time of 5.5 seconds per frame, and crystal-to-detector distance was 850 mm, at an X-ray wavelength of 1.03320 Å (Figure 3). Data were processed with HKL2000 (Otwinowski, 1997). Due to the crystal decay, only the first 70 frames were processed to yield a data set. Data collection and processing statistics are summarized in Table 1. Self-rotation function was calculated with the program Replace (Tong & Rossmann, 1990).

**Table 1.**
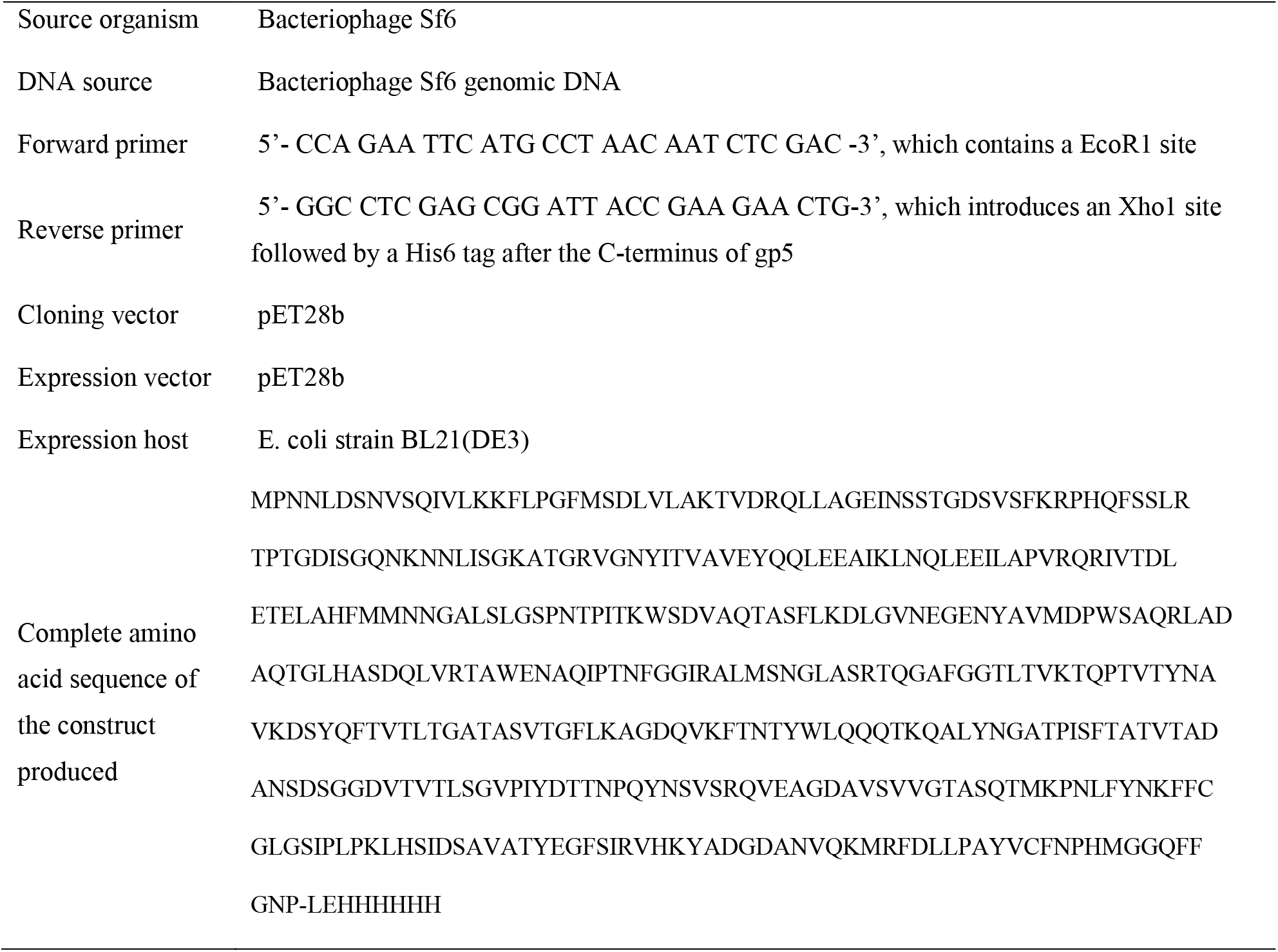
Macromolecule production information [Style: IUCr table caption; this style applies table numbering]

**Table 2.** Crystallization

|  |  |
| --- | --- |
| Method | Hanging drop vapour diffusion |
| Plate type | Hampton Research 24-well VDX plate |
| Temperature (K) | 293 |
| Protein concentration | 13.4mg/ml |
| Buffer composition of protein solution | 20mM Tris-HCl pH 8.0, 50mM NaCl and 1mM EDTA |
| Composition of reservoir solution | 5% polyethylene glycol 8000, 100 mM Tris-HCl pH 8.5 and 1M NaCl |
| Volume and ratio of drop | 1ul protein solution was mixed with 1ul well solution |
| Volume of reservoir | 500ul |

**Table 3.** Data collection and processing Values for the outer shell are given in parentheses.

|  |  |
| --- | --- |
| Diffraction source | APS GM/CA-CAT 23-ID-B |
| Wavelength (Å) | 1.03320 |
| Temperature (K) | 100 |
| Detector | MAR 300 CCD |
| Crystal-detector distance (mm) | 850 |
| Rotation range per image (°) | 1 |
| Total rotation range (°) | 131 |
| Exposure time per image (s) | 5.5 |
| Space group | *C2* |
| *a*, *b*, *c* (Å) | 973.33, 568.23, 565.57 |
| $\alpha$, $\beta$, $\gamma$ (°) | 90, 120.54, 90 |
| Mosaicity (°) | 0.54 |
| Resolution range (Å) | 50.00–5.90 (6.00–5.90) |
| Total No. of reflections | 346977 |
| No. of unique reflections | 212832 |
| Completeness (%) | 31.1 (12.9)# |
| Redundancy | 1.6 (1.1) |
| $\langle I/\sigma(I) \rangle$ | 3.9(1.2)† |
| $R_{\text{r.i.m.}}^{\ddagger}$ | 0.245 (1.423) |
| Overall *B* factor from Wilson plot (Å<sup>2</sup>) | 58.79 |

**Figure 1.**
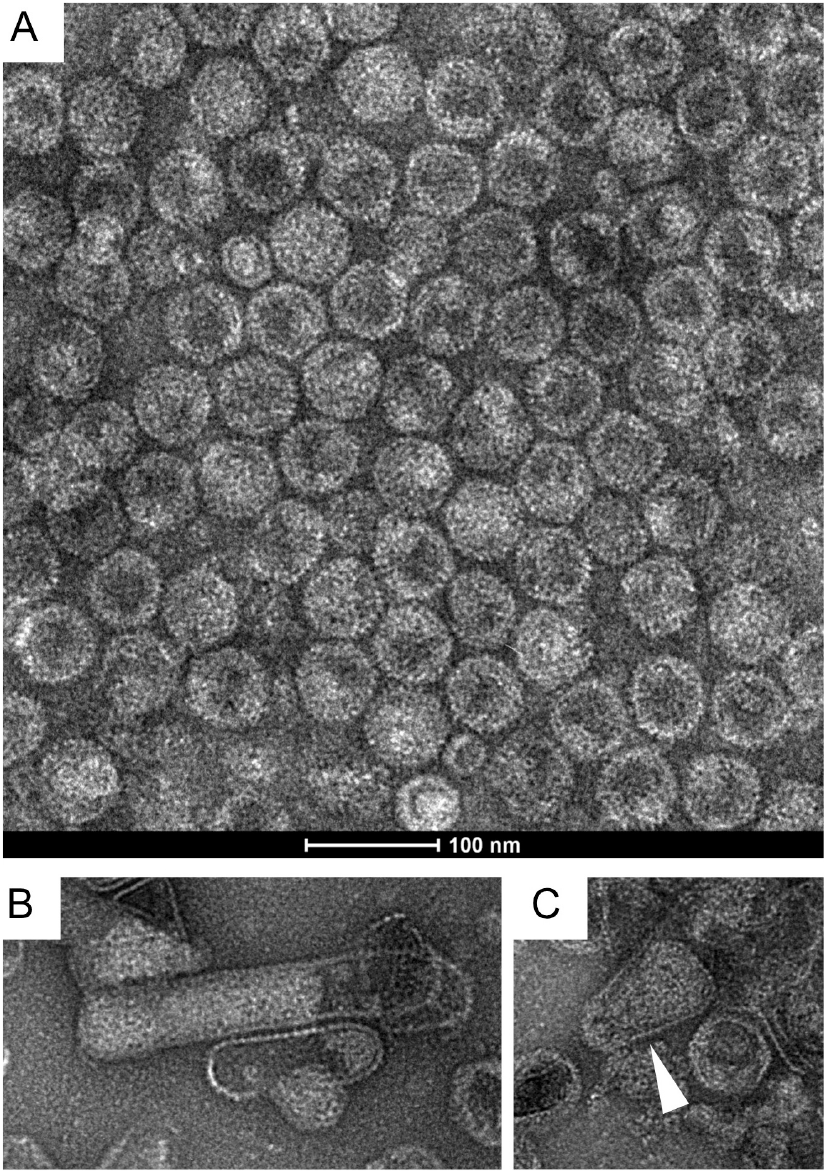
Electron micrographs of negatively stained gp5 PLPs. (A) Purified gp5 PLPs. (B) Elongated tube-like structure formed by gp5. (C) Cone-shaped structure formed by gp5 indicated with an arrowhead.

**Figure 2.**
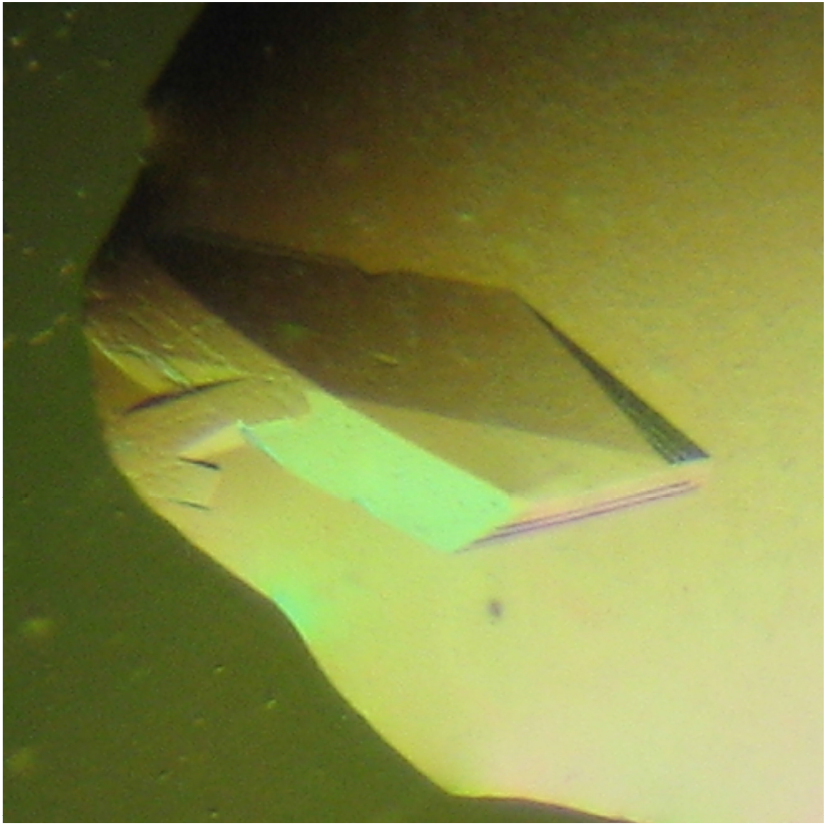
A representative crystal of gp5 PLPs. The crystal grew to the maximal size of 0.4×0.2×0.1 mm^3^ within two weeks.

**Figure 3.**
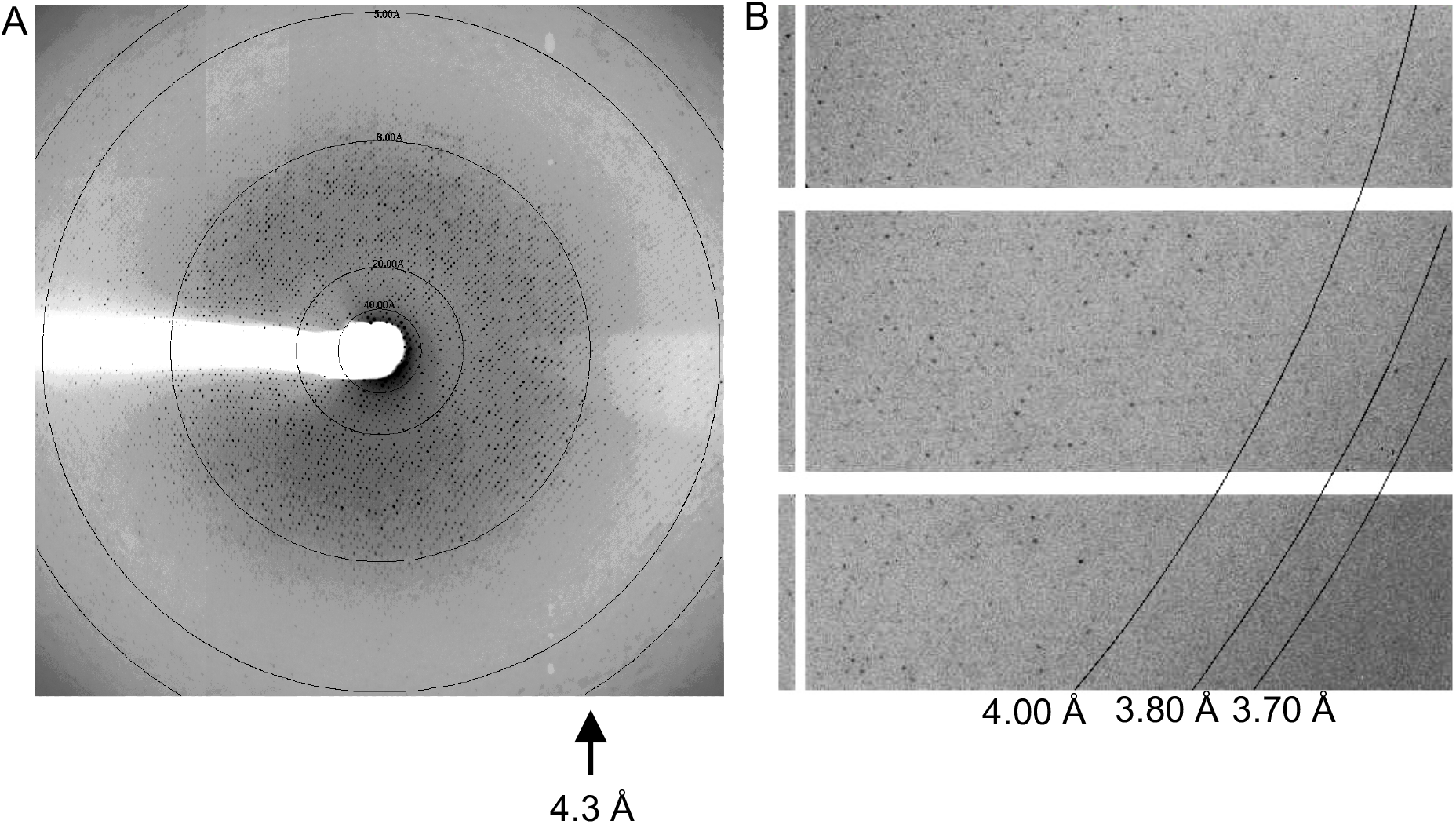
X-ray diffraction of gp5 PLP crystals. (A) A diffraction pattern collected with a MARmosaic CCD detector indicating diffraction beyond 4.3Å resolution. (B) An enlarged area in a diffraction pattern collected with a Pilatus3 6M detector. Note reflection spots beyond 4 Å resolution.

## 3. Results and discussion

### 3.1. Assembly of the gp5 PLPs

The Sf6 gp5 protein was over-expressed in *E. coli* at a yield of ∼15 mg purified protein per liter of culture medium. Negative staining electron microscopy showed that gp5 spontaneously assembled into isometric spherical particles with an overall diameter of 600 Å (Fig. 1 A), consistent with the appearance of procapsid. Other forms of particles were observed in the un-purified sample, including elongated tube-like structure resembling the phage P22 polyheads (Parent *et al*., 2010, Suhanovsky *et al*., 2010, Parent *et al*., 2007) (Fig. 1 B), and cone-shaped particles reminiscent of the human immune-deficiency virus capsid (Fig. 1 C) (Fig. 1 A). These data suggest that gp5 is capable of assembling into a variety of forms of particles. This is consistent with observations for phage P22 capsid protein, which forms various forms of particles in the absence of the scaffolding protein (Suhanovsky *et al*., 2010, Parent *et al*., 2007).

### 3.2. Crystallographic analysis

The purified gp5 procapsid-like particles were crystallized (Fig. 2). Crystals diffracted beyond 4.3 Å resolution (Fig. 3). Nevertheless, quick decay was observed for crystals at synchrotron radiation. A large number of crystals were screened in order to identify a single crystal that can survive X-ray exposure. A data set was collected and processed, which contains a total of 212,832 unique reflections in the resolution range 50.0-5.9 Å with a completeness of 31.1% and an *R*merge of 15.0% (Table 1). Crystals belong to the space group *C*2 with unit cell dimensions of a=973.33 Å, b=568.23 Å, c=565.57 Å, and β=120.54°. Self-rotation function clearly shows 532 symmetry (Fig. 4), confirming formation of icosahedral particles. There is a half of the *T*=7 icosahedral particle in the crystallographic asymmetric unit. This corresponds to a 82% solvent content and a Vm value of 7.5 Å^3^/Da (Matthews, 1968). The particle is situated in the crystal unit cell such that the icosahedral 2-fold axis coincides with the crystallographic *b* axis and the perpendicular icosahedral 2-fold axis coincides with the crystallographic *a* axis. Structure determination with the molecular replacement method using 30-fold non-crystallographic symmetry electron density averaging is underway.

**Figure 4.**
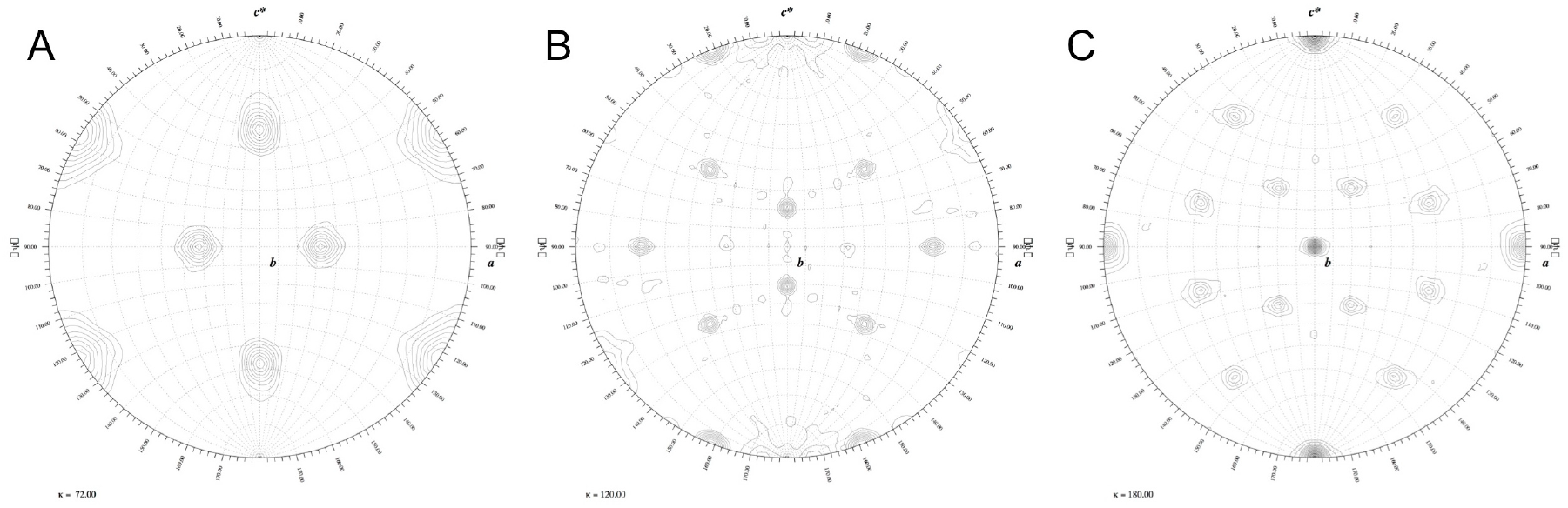
Self-rotation functions calculated of gp5 PLP crystal X-ray data for kappa of 72 degrees (A), 120 degrees (B) and 180 degrees (C), showing peaks for 5-, 3-, and 2-fold axes respectively.

## Acknowledgements

We thank the staff at the beamlines GM/CA-CAT 23-ID-D and 23-ID-B at the Advanced Photon Source for beamline access. This work was supported by the grant R01GM090010 from the National Institute of General Medical Sciences of the National Institutes of Health to L.T.

## References

Aksyuk, A. A., Bowman, V. D., Kaufmann, B., Fields, C., Klose, T., Holdaway, H. A., Fischetti, V. A. & Rossmann, M. G. (2012). Proc Natl Acad Sci U S A 109, 14001–14006.

Casjens, S., Winn-Stapley, D. A., Gilcrease, E. B., Morona, R., Kuhlewein, C., Chua, J. E., Manning, P. A., Inwood, W. & Clark, A. J. (2004). J Mol Biol 339, 379–394.

Dai, W., Fu, C., Raytcheva, D., Flanagan, J., Khant, H. A., Liu, X., Rochat, R. H., Haase-Pettingell, C., Piret, J., Ludtke, S. J., Nagayama, K., Schmid, M. F., King, J. A. & Chiu, W. (2013). Nature 502, 707–710.

Fokine, A., Leiman, P. G., Shneider, M. M., Ahvazi, B., Boeshans, K. M., Steven, A. C., Black, L. W., Mesyanzhinov, V. V. & Rossmann, M. G. (2005). Proc Natl Acad Sci U S A 102, 7163–7168.

Gemski, P., Jr., Koeltzow, D. E. & Formal, S. B. (1975). Infect Immun 11, 685–691.

Gertsman, I., Gan, L., Guttman, M., Lee, K., Speir, J. A., Duda, R. L., Hendrix, R. W., Komives, E. A. & Johnson, J. E. (2009). Nature 458, 646–650.

Guo, F., Liu, Z., Fang, P. A., Zhang, Q., Wright, E. T., Wu, W., Zhang, C., Vago, F., Ren, Y., Jakana, J., Chiu, W., Serwer, P. & Jiang, W. (2014). Proc Natl Acad Sci U S A 111, E4606–4614.

Lindberg, A. A., Wollin, R., Gemski, P. & Wohlhieter, J. A. (1978). J Virol 27, 38–44.

Matthews, B. W. (1968). J Mol Biol 33, 491–497.

Otwinowski, Z., and Minor, W. (1997). Methods Enzymol., pp. 307–326.

Parent, K. N., Gilcrease, E. B., Casjens, S. R. & Baker, T. S. (2012). Virology 427, 177–188.

Parent, K. N., Sinkovits, R. S., Suhanovsky, M. M., Teschke, C. M., Egelman, E. H. & Baker, T. S. (2010). Phys Biol 7, 045004.

Parent, K. N., Suhanovsky, M. M. & Teschke, C. M. (2007). Mol Microbiol 65, 1300–1310.

Parent, K. N., Tang, J., Cardone, G., Gilcrease, E. B., Janssen, M. E., Olson, N. H., Casjens, S. R. & Baker, T. S. (2014). Virology 464–465, 55-66.

Rizzo, A. A., Suhanovsky, M. M., Baker, M. L., Fraser, L. C., Jones, L. M., Rempel, D. L., Gross, M. L., Chiu, W., Alexandrescu, A. T. & Teschke, C. M. (2014). Structure 22, 830–841.

Suhanovsky, M. M., Parent, K. N., Dunn, S. E., Baker, T. S. & Teschke, C. M. (2010). Mol Microbiol 77, 1568–1582.

Suhanovsky, M. M. & Teschke, C. M. (2015). Virology 479–480, 487-497.

Sun, S., Kondabagil, K., Draper, B., Alam, T. I., Bowman, V. D., Zhang, Z., Hegde, S., Fokine, A., Rossmann, M. G. & Rao, V. B. (2008). Cell 135, 1251–1262.

Tong, L. A. & Rossmann, M. G. (1990). Acta Crystallogr A 46 (Pt 10), 783–792.

van Heel, M., Gowen, B., Matadeen, R., Orlova, E. V., Finn, R., Pape, T., Cohen, D., Stark, H., Schmidt, R., Schatz, M. & Patwardhan, A. (2000). Q Rev Biophys 33, 307–369.

Veesler, D. & Johnson, J. E. (2012). Annual review of biophysics 41, 473–496.

Wikoff, W. R., Liljas, L., Duda, R. L., Tsuruta, H., Hendrix, R. W. & Johnson, J. E. (2000). Science 289, 2129–2133.

Zhao, H., Christensen, T. E., Kamau, Y. N. & Tang, L. (2013). Proc Natl Acad Sci U S A 110, 8075–8080.

Zhou, Z. H., Hui, W. H., Shah, S., Jih, J., O’Connor, C. M., Sherman, M. B., Kedes, D. H. & Schein, S. (2014). Structure 22, 1385–1398.

